# Is thermal aptitude a pivotal driver in the establishment of recent *Puccinia striiformis* f. sp. *tritici* lineages in Europe?

**DOI:** 10.1101/2023.08.11.552914

**Authors:** Kevin JG Meyer, Marc Leconte, Tiphaine Vidal, Henriette Goyeau, Frédéric Suffert

## Abstract

In the context of global warming, it is crucial to focus on the effects of temperature on the emergence of new lineages of endemic pathogen species, such as *Puccinia striiformis* f. sp. *tritici* (*Pst*) the causal agent of yellow rust on wheat. We characterized the thermal aptitude of representative isolates from the most recent common European *Pst* races. We assessed two key aggressiveness components – infection efficiency (IE) and latency period (LP) – under warm and cold thermal regimes, comparing 10 *Pst* isolates collected from 2010 to 2020 with three “old” reference isolates. The significant differences observed suggest that this species has the potential to adapt to temperature changes, but that such adaptation probably did not drive the establishment of the most recent races and the dominant ‘Warrior’ and ‘Warrior(-)’ they succeeded. These races display “generalist” behavior with respect to temperature, with ‘Warrior(-)’ showing no more aggressiveness than the races replaced since the 1990s. The differences in competitive success between emerging *Pst* lineages are probably due to the deployment of resistance genes in wheat and the advantages of new forms of virulence emerging independently of thermal adaptability. However, variations in thermal adaptability for both aggressiveness components suggested an impact of geographic origin within the ‘Warrior’ and ‘Warrior(-)’ races, as previously reported for the “old” reference isolates. Furthermore, the independence of thermal adaptability established for IE and LP implies that the effects of temperature may depend on the stage of the epidemic (early or late), potentially modifying seasonal dynamics.

## Introduction

Plant pathogens can adapt to both biotic changes, such as alterations to the varietal landscape driven by the deployment of resistance genes (McDonald and Linde, 2002), and abiotic changes, such as changes in temperature (Helfer *et al.,* 2014). Thermal adaptation is considered particularly important given that global mean surface temperature increased by 1.1°C from 1850-1900 to 2011-2020 (Gulev *et al*., 2021), and global warming is continuing, with a likely 1.5°C increase by the early 2030s (Lee *et al*., 2021). Water cycles around the world are also displaying alterations (Douville *et al*., 2021). In Europe, the frequency of extreme weather conditions has also increased in recent decades, with an impact on human activities and agricultural resources (Andrade *et al*., 2012).

Studies of adaptation to biotic changes have focused on compatible gene-for-gene interactions between plant pathogens and their hosts (Figueroa *et al*., 2020), to characterize the virulence profiles defining races in the case of wheat rusts, with two pathogenic isolates considered to belong to the same race if they have the same combination of virulence genes. In *Puccinia striiformis* f. sp. *tritici* (*Pst*), the causal agent of yellow rust, changes in these profiles are closely linked to the corresponding resistances in wheat populations (Thrall & Burdon, 2003), accounting for the emergence of major *Pst* races, and generally leading to the evolution of populations being interpreted as a change in virulence profiles. Populations dominated by a limited number of races are known to be selected by the wheat varieties deployed in the landscape (de Vallavieille *et al*., 2012), and sexual reproduction generates new opportunistic clonal lineages carrying new combinations of virulence genes (Figueora *et al*., 2020; Hovmøller *et al*., 2016). Sexual reproduction in Central Asia (Ali *et al*., 2017) led to the recent replacement of major *Pst* races in Europe through a suspected incursion of exotic isolates from this part of the world, resulting in the establishment of the Warrior race across the continent in 2011 (Hovmøller *et al*., 2016; Hubbard *et al*., 2015). The European RUSTWATCH project has recently identified a new ‘Warrior(-)’ race from a different genetic group replacing ‘Warrior’ (https://agro.au.dk/forskning/internationale-platforme/wheatrust/yellow-rust-tools-maps-and-charts/races-changes-across-years). It has also been shown, in wheat leaf rust due to *Puccinia triticina,* that the distribution of resistance genes in the landscape is insufficient in itself to account for all the variation in plant pathogen race-host cultivar associations (Fontyn *et al*., 2022). Other selection factors relating to quantitative components of plant-pathogen interactions must, therefore, be considered.

Responses to abiotic changes have been observed in both host plants and plant pathogens (Garett *et al*., 2006; Chen *et al*., 2017), with adult plant resistance directly influenced by temperature (Rodriguez-Algaba *et al*., 2019), and a possible decrease in the resistance of cultivated plants with changing climatic conditions, such as the increasing dryness of the climate (Garett *et al*. 2006). The natural ranges of rust species have also expanded, with an incursion of *Pst* into South Africa reported following changes in rainfall patterns (Boshoff *et al*., 2002), and the possible establishment of *Puccinia graminis* f. sp. *tritici* (*Pgt*) and *Pst* in new regions, given their higher rates of survival in milder winters (Prank *et al*., 2019; Ma *et al.,* 2015; Novotná *et al*., 2017), resulting in a higher risk of crop diseases in a context of climate change (Juroszek *et al*., 2020). Temperature is an abiotic factor that is easy to monitor, and considered to drive changes in pathogen biology (Chen *et al*., 2017): thermal effects have been demonstrated on the sexual and asexual parts of fungal reproductive cycles (McDonald and Linde, 2002), including the formation of larger numbers of telia and teliospores at high temperature in *Pst* (Chen *et al*., 2021).

One of the most widely monitored features of the adaptation of plant pathogens is their aggressiveness, which determines the rate at which a given disease intensity is reached and can therefore be used as a proxy for the total amount of damage to plants in the field (Andrivon *et al*. 2007; Lannou 2012). The assessment of aggressiveness is intrinsically complex because this characteristic is related to various life-history traits of the pathogen that must be measured during the host-pathogen interaction. The most widely assessed traits for rust pathogens are infection efficiency, latency period and sporulation capacity (Pariaud *et al*., 2009; Milus *et al*., 2006; Fontyn *et al*., 2022). Infection efficiency is the proportion of spores able to cause a new infection when deposited on compatible host-plant tissues. Latency period is the time between the deposition of a spore and the appearance of most of the sporulating structures. Sporulation capacity is the number of spores produced per individual sporulating structure and per unit time. Changes in one or several of these aggressiveness components may cause population shifts, even if the virulence profile is not affected (Milus *et al*., 2009). As stated above, the distribution of resistance genes in the landscape is insufficient in itself to account for all the variation in plant pathogen race-host cultivar associations, and components of aggressiveness have been shown to be important drivers of the evolution of pathogen populations in *P. triticina* (Fontyn *et al*., 2023). Conversely, a decrease in aggressiveness can be compensated by a gain in virulence to overcome a resistance gene carried by one or more varieties, particularly if they are deployed over large areas. Such a trade-off between virulence and aggressiveness is known to occur in some plant pathogens, and pathogens from more susceptible host populations tend to evolve towards greater aggressiveness and a loss of unnecessary virulence genes (Thrall & Burdon, 2003), highlighting the importance and regulation of aggressiveness in the face of changing biotic conditions.

Evidence for thermal adaptation has been obtained in Euro-Mediterranean populations of the wheat pathogen *Zymoseptoria tritici* at contrasting spatiotemporal scales. Geographic variations of thermal responses were found across contrasting climatic areas, together with local changes due to seasonal patterns over single wheat-growing seasons (Suffert *et al*., 2016; Boixel *et al*., 2022). Pathogen populations collected from susceptible cultivars were found to be more aggressive at lower temperatures, whereas no effect of temperature was observed on moderately resistant cultivars (Chen *et al*., 2017), implying that interactions between host resistance and temperature are key drivers of pathogen evolution. Recent populations of *Pst* in the United States have been shown to have an optimum temperature of about 18°C for spore production and a shorter latency period than the populations from before the 2000s, which had an optimum of 13°C to 16°C, revealing an adaptation of pathogen populations to higher temperatures (Milus *et al*., 2006). *Pst* has generally been considered mostly a mild/cool-climate pathogen limited by warmer temperatures (e.g., Dennis, 1987). Its adaptation to warmer temperatures is therefore likely to serve as a very potent lever for invasion of new areas (Vidal *et al.,* 2022). In Europe, isolates of the ‘Warrior’ race have been shown to have generalist thermal behavior, with similar performances under different thermal regimes, resulting in an intermediate capacity to tolerate a warming of the climate (de Vallavieille-Pope *et al*., 2018). The structure of the French *Pst* population before 2004 has revealed local thermal adaptation, with a significant pathogen geographic origin (southern vs. northern France) × temperature interaction for urediniospore germination rate and infection efficiency (Mboup *et al*., 2012). Going beyond the demonstration of such local adaptation to temperature for some particular *Pst* races, we need to understand whether the recent evolution of *Pst* populations at European scale is related to variations in the performance of newly identified races in response to increasing temperatures and, if so, to consider the geographic origin of these races.

We tested the hypothesis of an adaptation of several recent European races of *Pst* to warm temperatures by comparing the infection efficiency and latency period of isolates representative of the most common European races collected over the last decade (2010-2020) with those of “old” reference isolates collected before the 1990s, under different thermal regimes. We paid particular attention to ‘Warrior’ (PstS7), and ‘Warrior(-)’ (PstS10), which has a more limited virulence profile and has been gradually replacing ‘Warrior’ in Europe since 2014.

## Materials and methods

### Overall experimental design

Seedlings of bread wheat cv ‘Michigan Amber’, which is known to be susceptible to yellow rust, were inoculated with 13 *Pst* isolates from nine different races, spanning different time periods and originating from different geographic areas in Western Europe. Two aggressiveness components – infection efficiency (IE) and latency period (LP) – were measured under different temperature conditions, as described by Vallavieille Pope *et al*. (2018). For IE measurements, seedlings were subjected to four different thermal regimes (5°C, 10°C, 15°C, and 20°C; Figure 1) during the first 24 hours post-inoculation (hpi), when the spores were germinating on the leaf surface. All seedlings were subsequently incubated in identical conditions (a 16 h light/8 h dark photoperiod, with temperatures of 20°C during light periods and 15°C during periods in the dark). The first symptoms of infection were observed six to seven days post-inoculation (dpi). For LP measurements, seedlings were kept in the same optimal conditions during the first 24 hpi after inoculation, i.e. ‘incubation period’ (8.5°C; de Vallavieille Pope *et al*., 2018) and were then subjected to one of two thermal regimes over the following 20 days: 16 h light/8 h dark photoperiod with temperatures of 25°C during light periods and 16°C during dark periods, mimicking a ‘warm regime’, or a 16 h light/8 h dark photoperiod with temperatures of 15°C during light periods and 10°C during dark periods, mimicking a ‘cold regime’. Symptoms were observed 19 to 21 dpi. Each experiment was conducted twice.

**Figure 1.**
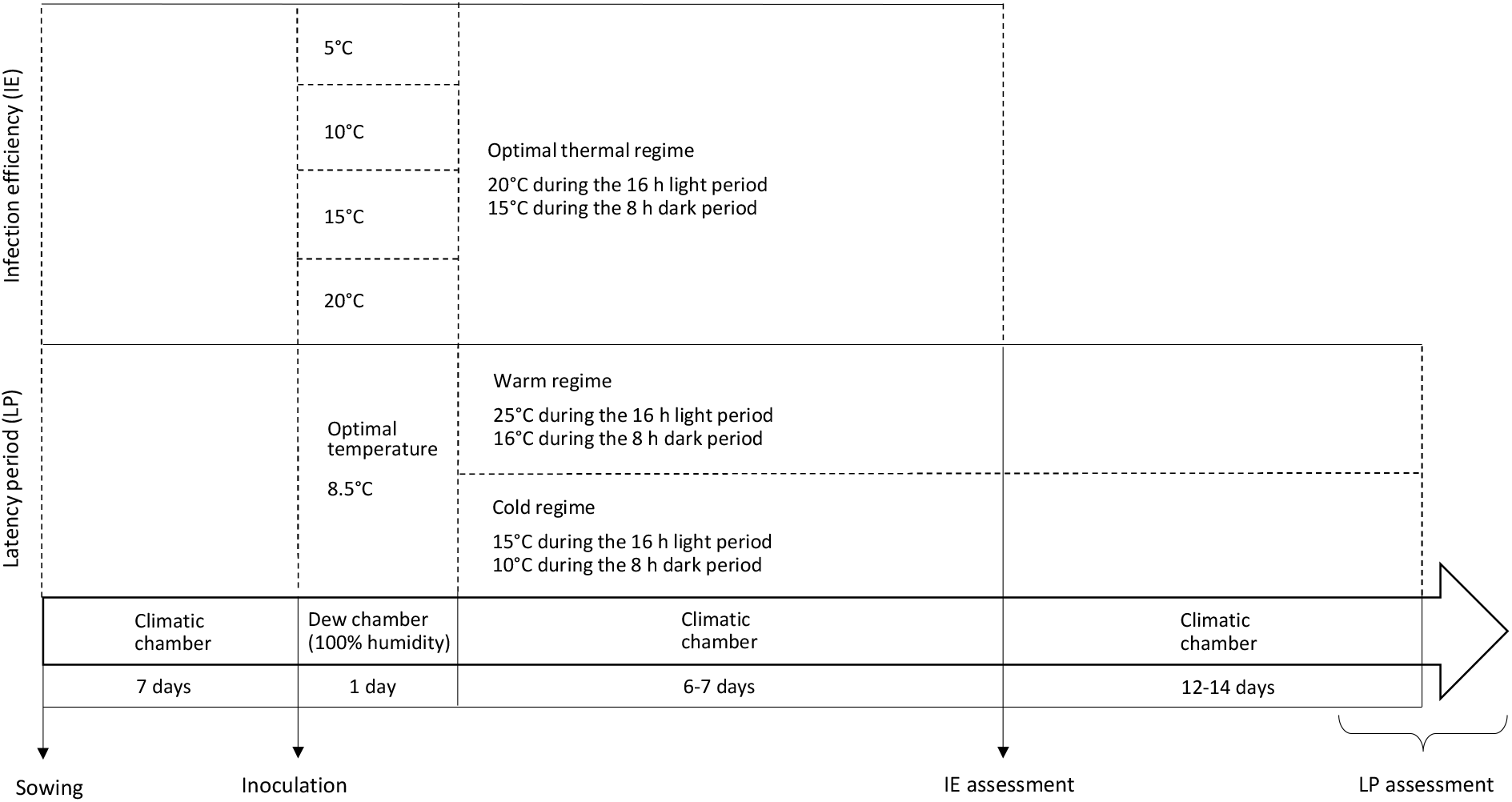
Overview of the experimental protocol for evaluating the infection efficiency (IE) and latency period (LP) of *Puccinia striiformis* f. sp. *tritici* isolates under different thermal regimes.

### Plant material

We sowed 15 seeds of the bread wheat cv ‘Michigan Amber’ in square pots (7 x 7 x 8 cm). Wheat seedlings were grown in a climatic chamber to the two-leaf stage at 20°C during the 16 h light period and 15°C during the 8 h dark period. Plants were exposed to artificial light for 24 hours on the day before inoculation, as increasing duration of illumination increases IE values (de Vallavielle-Pope *et al*., 2012), facilitating comparisons. Immediately before inoculation, 10 homogeneous plants per pot were selected, and their second leaves were cut off.

### Fungal material

The *Pst* isolates studied were chosen so as to best represent the recent European population (period 2010-2020). We characterized the thermal aptitude of these isolates relative to the three reference isolates, by measuring IE and LP as aggressiveness traits (Table 1).

**Table 1.**
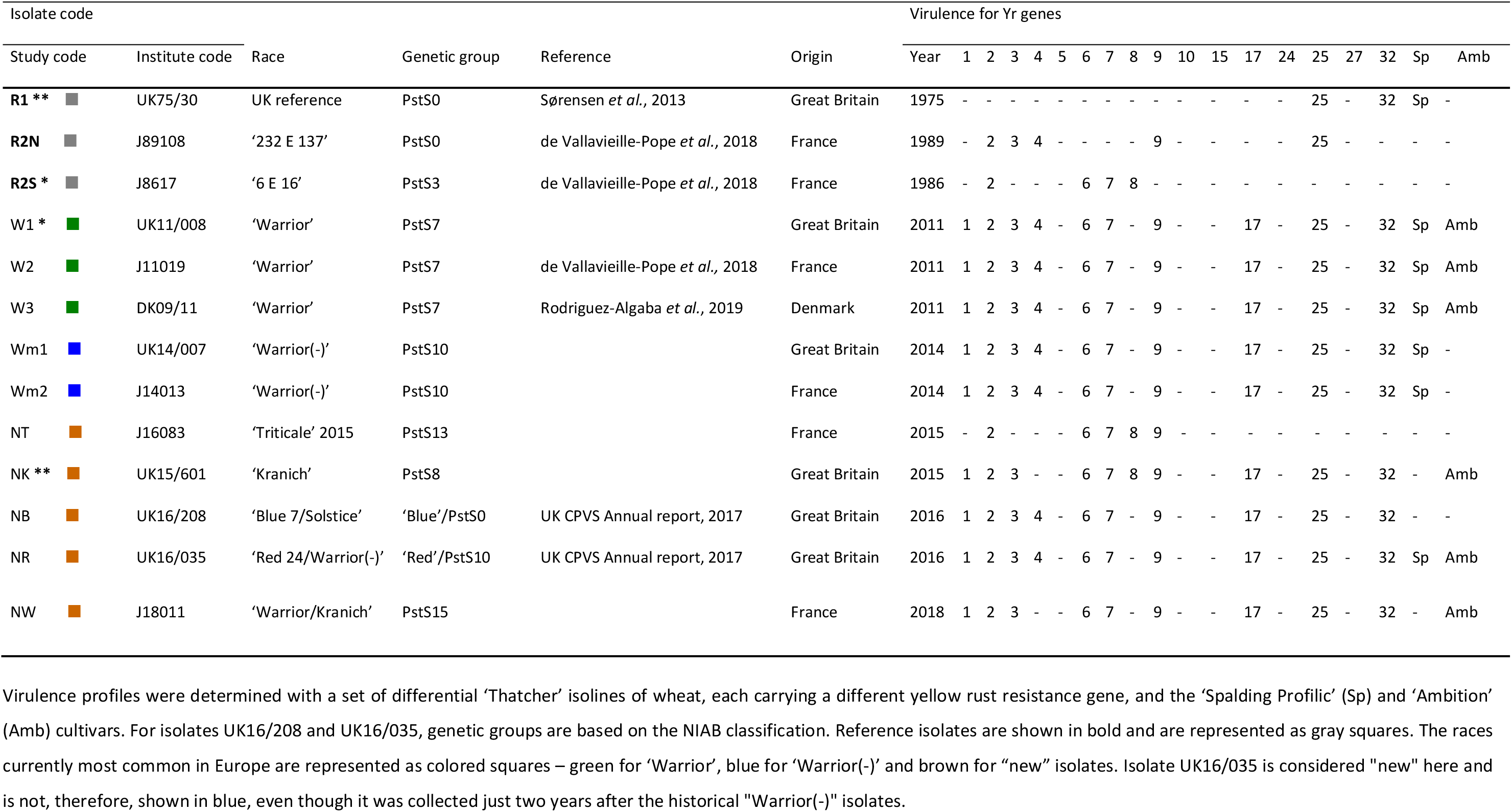
List of the 13 *Puccinia striiformis* f. sp. *tritici* isolates for which infection efficiency (IE) and latency period (LP) were assessed

| Isolate code |  |  |  |  |  | Virulence for Yr genes |  |  |  |  |  |  |  |  |  |  |  |  |  |  |  |  |  |  |
| --- | --- | --- | --- | --- | --- | --- | --- | --- | --- | --- | --- | --- | --- | --- | --- | --- | --- | --- | --- | --- | --- | --- | --- | --- |
| Study code | Institute code | Race | Genetic group | Reference | Origin | Year | 1 | 2 | 3 | 4 | 5 | 6 | 7 | 8 | 9 | 10 | 15 | 17 | 24 | 25 | 27 | 32 | Sp | Amb |
| **R1 \*\*** | UK75/30 | UK reference | PstS0 | *Sørensen et al., 2013* | Great Britain | 1975 | - | - | - | - | - | - | - | - | - | - | - | - | - | 25 | - | 32 | Sp | - |
| **R2N** | J89108 | '232 E 137' | PstS0 | *de Vallavieille-Pope et al., 2018* | France | 1989 | - | 2 | 3 | 4 | - | - | - | - | 9 | - | - | - | - | 25 | - | - | - | - |
| **R2S \*** | J8617 | '6 E 16' | PstS3 | *de Vallavieille-Pope et al., 2018* | France | 1986 | - | 2 | - | - | - | 6 | 7 | 8 | - | - | - | - | - | - | - | - | - | - |
| W1 * | UK11/008 | 'Warrior' | PstS7 |  | Great Britain | 2011 | 1 | 2 | 3 | 4 | - | 6 | 7 | - | 9 | - | - | 17 | - | 25 | - | 32 | Sp | Amb |
| W2 | J11019 | 'Warrior' | PstS7 | *de Vallavieille-Pope et al., 2018* | France | 2011 | 1 | 2 | 3 | 4 | - | 6 | 7 | - | 9 | - | - | 17 | - | 25 | - | 32 | Sp | Amb |
| W3 | DK09/11 | 'Warrior' | PstS7 | *Rodriguez-Algaba et al., 2019* | Denmark | 2011 | 1 | 2 | 3 | 4 | - | 6 | 7 | - | 9 | - | - | 17 | - | 25 | - | 32 | Sp | Amb |
| Wm1 | UK14/007 | 'Warrior(-)' | PstS10 |  | Great Britain | 2014 | 1 | 2 | 3 | 4 | - | 6 | 7 | - | 9 | - | - | 17 | - | 25 | - | 32 | Sp | - |
| Wm2 | J14013 | 'Warrior(-)' | PstS10 |  | France | 2014 | 1 | 2 | 3 | 4 | - | 6 | 7 | - | 9 | - | - | 17 | - | 25 | - | 32 | Sp | - |
| NT | J16083 | 'Triticale' 2015 | PstS13 |  | France | 2015 | - | 2 | - | - | - | 6 | 7 | 8 | 9 | - | - | - | - | - | - | - | - | - |
| NK ** | UK15/601 | 'Kranich' | PstS8 |  | Great Britain | 2015 | 1 | 2 | 3 | - | - | 6 | 7 | 8 | 9 | - | - | 17 | - | 25 | - | 32 | - | Amb |
| NB | UK16/208 | 'Blue 7/Solstice' | 'Blue'/PstS0 | UK CPVS Annual report, 2017 | Great Britain | 2016 | 1 | 2 | 3 | 4 | - | 6 | 7 | - | 9 | - | - | 17 | - | 25 | - | 32 | - | - |
| NR | UK16/035 | 'Red 24/Warrior(-)' | 'Red'/PstS10 | UK CPVS Annual report, 2017 | Great Britain | 2016 | 1 | 2 | 3 | 4 | - | 6 | 7 | - | 9 | - | - | 17 | - | 25 | - | 32 | Sp | Amb |
| NW | J18011 | 'Warrior/Kranich' | PstS15 |  | France | 2018 | 1 | 2 | 3 | - | - | 6 | 7 | - | 9 | - | - | 17 | - | 25 | - | 32 | - | Amb |

Virulence profiles were determined with a set of differential 'Thatcher' isolines of wheat, each carrying a different yellow rust resistance gene, and the 'Spalding Profilic' (Sp) and 'Ambition' (Amb) cultivars. For isolates UK16/208 and UK16/035, genetic groups are based on the NIAB classification. Reference isolates are shown in bold and are represented as gray squares. The races currently most common in Europe are represented as colored squares – green for 'Warrior', blue for 'Warrior(-)' and brown for "new" isolates. Isolate UK16/035 is considered "new" here and is not, therefore, shown in blue, even though it was collected just two years after the historical "Warrior(-)" isolates.

Thirteen isolates were selected from the mycotheques of European research laboratories working on yellow rust. Three “old” reference isolates collected before the incursion of Warrior were chosen: the French ‘6 E 16’ race as a ‘southern reference’ (R2S; collected in 1986), the French ‘232 E 137’ race as a ‘northern reference’ (R2N; collected in 1989) and the ‘UK75/30’ race from UK (R1; collected in 1975), which carries very few virulence genes. A Danish isolate from the common ‘Warrior’ race group collected in 2011 was also used. Three common recent French races were included in the set (‘Warrior’, ‘Warrior(-)’ and ‘Triticale’ from 2011, 2014 and 2015, respectively) along with a recent French isolate from the ‘Warrior/Kranich’ race. Five isolates from the UK were also studied: three isolates from the ‘Warrior’, ‘Warrior(-)’ from 2014 and ‘Kranich’ races, and two isolates from the recent races ‘Blue 7/Solstice’ and ‘Red 24/Warrior(-)’, according to the NIAB classification (UK CPVS Annual report, 2017). The ‘Red 24/Warrior(-)’ and ‘Blue 7/Solstice’ races are considered to be similar to ‘Warrior(-)’ (Pst10 genetic group) and ‘Solstice’ (Pst0 genetic group), corresponding to European pre-‘Warrior’ and ‘Warrior(-)’ races (Table 1). Together, these races are representative of the changes and incursions that have occurred in Europe in the last decade, given that the ‘Warrior’ (PstS7) and ‘Warrior(-)’ races expanded rapidly in European wheat-growing areas in 2011 and 2014, respectively (Ali *et al*., 2017).

### Infection efficiency assessment

For each *Pst* isolate, eight pots of 10 ‘Michigan Amber’ seedlings were inoculated with 1 mg of urediniospores diluted in 1 mL Novec oil, corresponding to 110 spores.cm^-2^, as described by Vallavieille-Pope *et al*. (2018). The pots were placed in a dew chamber (100% humidity) at one of the tested temperatures (5°C, 10°C, 15°C, or 20°C) for 24 hours (Figure 2A). The seedlings were then transferred to a climatic chamber under a single thermal regime considered optimal for the incubation of *Pst* (16 h light: 8 h dark photoperiod, with temperatures of 20°C during the light periods and 15°C during the dark periods). The number of chlorotic spots on the leaf was then determined in a defined area (an area of 4-5 cm^2^ in the middle of each leaf) to assess IE, which was calculated as the ratio of the mean number of chlorotic spots on the leaf 6 to 7 dpi divided by the estimated number of spores deposited (de Vallavieille-Pope *et al*., 2018).

**Figure 2.**
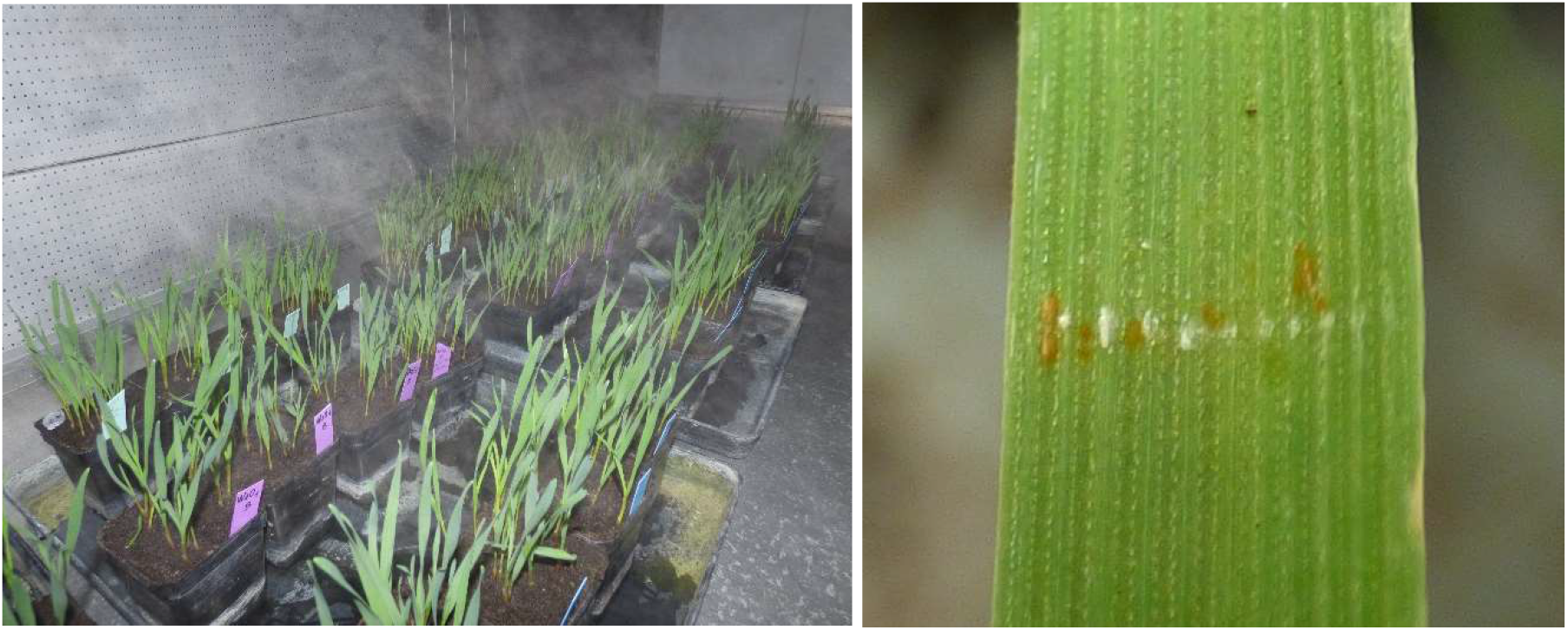
A. Seedlings placed in the dew chamber just after inoculation for the assessment of latency period (LP). B. Sporulating area observed at 8 dpi on the upper surface of the leaf for the assessment of infection efficiency (IE) after the application of *Puccinia striiformis* f. sp. *tritici* urediniospores with the edge of a plastic label covered with a mixture of 1 mg of spores in 25 mg of talcum powder.

### Latency period

For each *Pst* isolate, we inoculated eight pots of 10 ‘Michigan Amber’ seedlings with 1 mg of urediniospores mixed with 25 mg of talcum powder. The inoculum was applied on the upper face of the leaf by gently pressing the edge of a plastic label (1 mm thick) coated with the talcum powder/urediniospore mixture onto the leaf to deposit a narrow band of spores (Sørensen *et al*., 2013). The seedlings were then placed in a dew chamber (100% humidity) at 8.5°C for 24 hours, before being transferred to a climatic chamber under either the cold regime (15°C during the 16 h light period and 10°C during the 8 h dark period) or the warm regime (25°C during the 16 h light period and 16°C during the 8 h dark period) for 20 days. The number of seedlings with sporulating lesions was counted daily from day 8 to 20 dpi (Figure 2B), making it possible to evaluate the LP as the number of hpi to the appearance of sporulating lesions on half of the inoculated seedlings (de Vallavieille-Pope *et al*., 2018).

### Statistical analysis

All the analyses were performed with R software (R Core Team, 2022) version 4.0.3. Shapiro-Wilk tests (“shapiro.test” function) showed that IE and LP were not normally distributed. Non-parametric tests were therefore used to compare means: Wilcoxon tests (“wilcox.test” function) for two-level factors and Kruskal-Wallis tests (“kruskal” function; “agricolae*”* package, de Mendirburu and Yassen, 2020) for factors with 3 or more levels. Multivariate data analysis was performed by principal component analysis (“PCA” function from the “FactoMineR” package; Lê *et al*., 2008).

## Results

### Infection efficiency (IE)

IE was highest at 5°C, for all isolates, ranging from 10.5% for W1 to 26.0% for NT (Table S1), consistent with the published thermal optima, and IE was very low (<0.7%) at 20°C. Significant differences (*p*-value <0.05, Kruskal-Wallis test) were found between isolates at all temperatures tested, but no clearly homogeneous groups emerged (Figure 3).

**Figure 3.**
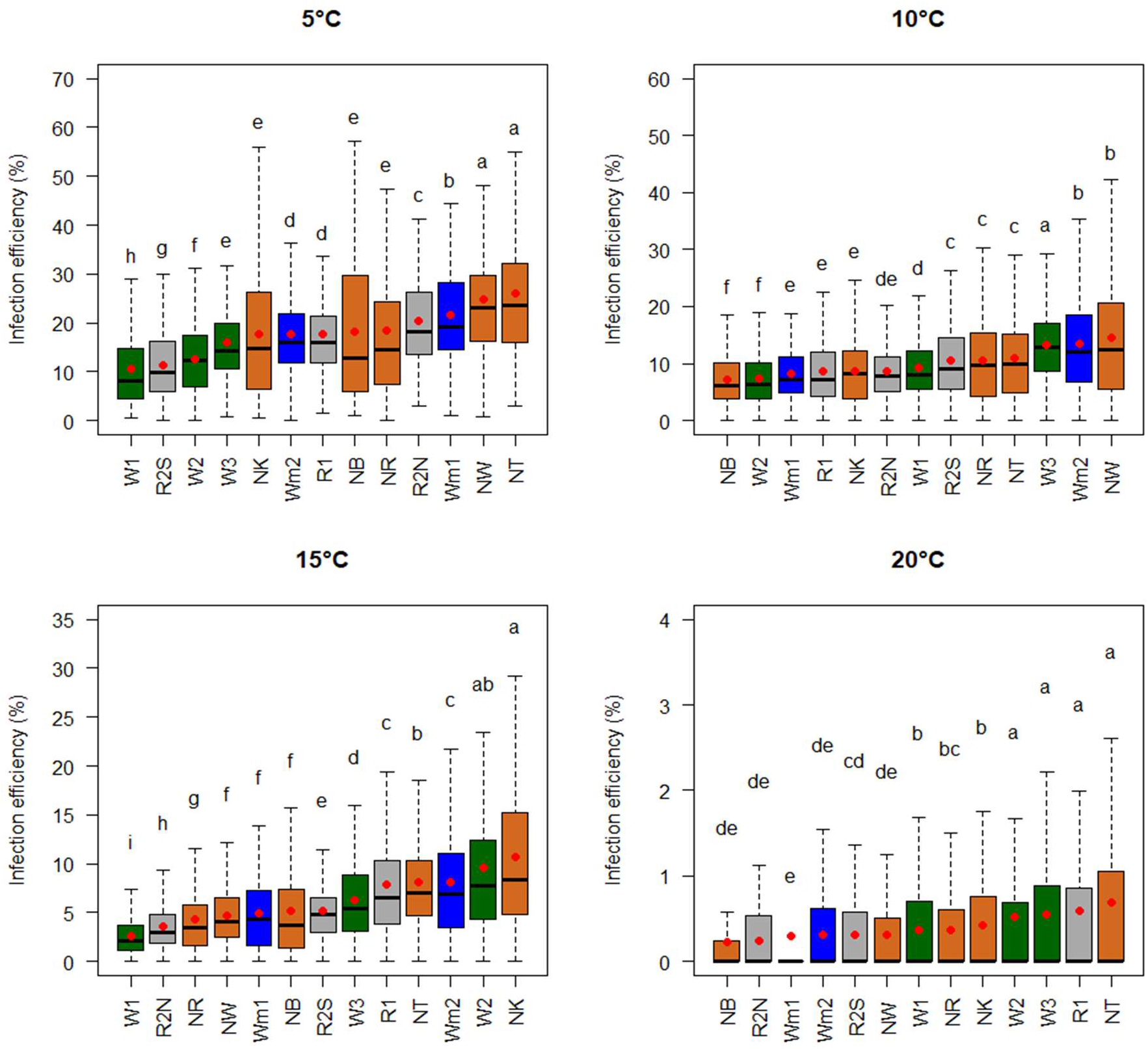
Mean infection efficiency (IE) and variability of IE for the 13 *Puccinia striiformis* f. sp. *tritici* isolates incubated at 5°C, 10°C, 15°C and 20°C during the first 24 h after inoculation. Letters indicate significant differences (*p*-value <0.05; Kruskal-Wallis tests). Red dots indicate the mean values. The races currently most common in Europe are indicated by the color of the box: green for ‘Warrior’, blue for ‘Warrior(-)’ and brown for the “new” isolates. The old reference isolates are represented by gray boxes.

The four isolates with the lowest IE at 5°C were the three representatives of the ‘Warrior’ race (W1, W2 and W3) and the southern reference race ‘232 E 137’ (R2S) (Figure 3). The four isolates with the highest IE at 5°C corresponded to the ‘Triticale’ and ‘Warrior/Kranich’ races (NT and NW) and, to a lesser extent, the French northern reference isolate (R2N) and the ‘Warrior(-)’ isolates from the UK (Wm1). Other isolates displayed intermediate behavior.

The isolates within each group behaved in various ways (Figure 4). For the old reference group, the French northern reference isolate, R2N, had a significantly higher IE at 5°C (20.3%) than the southern reference isolate, R2S (11.4%), whereas IE at 10°C and 15°C was higher for R2S (10.6% and 5.1%, respectively) than for R2N (8.7% and 3.6%, respectively). Recent isolates also displayed a diversity of behaviors within the same race. This heterogeneity was particularly pronounced around the optimal temperature in comparisons of IE between two isolates within the ‘Warrior’ or ‘Warrior(-)’ race, depending on temperature (5°C, 10°C or 15°C) Nevertheless, in several cases, the isolate with the highest IE at low temperature (5°C or 10°C) was among those with the lowest IE at higher temperature (10°C or 15°C), as shown in Figure 4. The ‘Warrior(-)’ isolate from the UK (Wm1) performed better than the ‘Warrior(-)’ isolate from France (Wm2) at 5°C (21.7% vs. 17.6%), but less well at 10°C and 15°C (8.3% vs. 13.4% and 4.9% vs. 8.2%). Similarly, the ‘Warrior’ isolates from Denmark (W3) and from the UK (W1) performed better than the ‘Warrior’ isolate from France (W2) at 10°C (13.2% and 9.2%, respectively, vs. 7.3%), but less well at 15°C (6.2% and 2.6%, respectively, vs. 9.6%). Furthermore, the Danish and British ‘Warrior’ isolates may be considered to behave similarly, in that there was no “inversion” of the ranking of their thermal performances from 5°C to 20°C. The French isolate was the most distinctive of the three ‘Warrior’ isolates, displaying differences from the other two.

**Figure 4.**
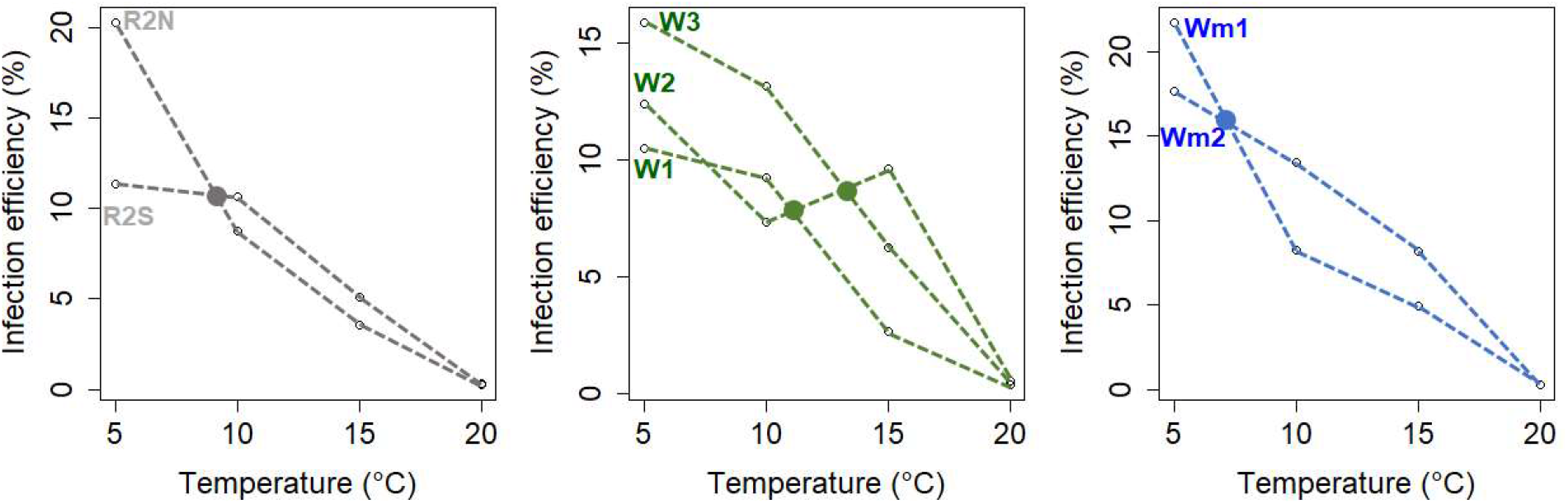
Thermal plasticity of infection efficiency (IE) for 7 *Puccinia striiformis* f. sp. *tritici* isolates from the reference group (R2N, R2S; gray lines), the ‘Warrior’ group (W1, W2, W3; green lines) and the ‘Warrior(-)’ group (Wm1, Wm2; blue lines), from 5°C to 20°C. The main crossovers of the thermal performance lines, marked with a colored circle, suggest an effect of geographic origin within each group (southern vs. northern; see Table 1 for this origin and Figure 3 for the significance of differences between isolates for each temperature).

### Latency period (LP)

Significant differences in LP were established for all isolates, for both the cold (15°C-10°C) and warm (25°C-16°C) regimes (Figure 5). Mean LP ranged from 203 hpi (NK) to 248 hpi (R1) under the cold regime and from 215 hpi (NW) to 330 hpi (R1) under the warm regime (Figure 5). Most of the isolates had a shorter LP under the cold regime than under the warm regime (Figure 6), consistent with the cold regime temperatures being closer to the optimum for *Pst*, as shown in previous studies (Mboup *et al*., 2012; de Vallavieille-Pope *et al*., in 2018). The northern French reference isolate (R2N) and the UK reference isolate (R1) had longer LPs than the most recent races, under both regimes (Figure 5), highlighting an overall increase in the aggressiveness of new European populations of *Pst*, of which LP is an important component. The southern French reference isolate (R2S) had a longer LP than the other isolates under the cold regime, and a shorter LP under the warm regime.

**Figure 5.**
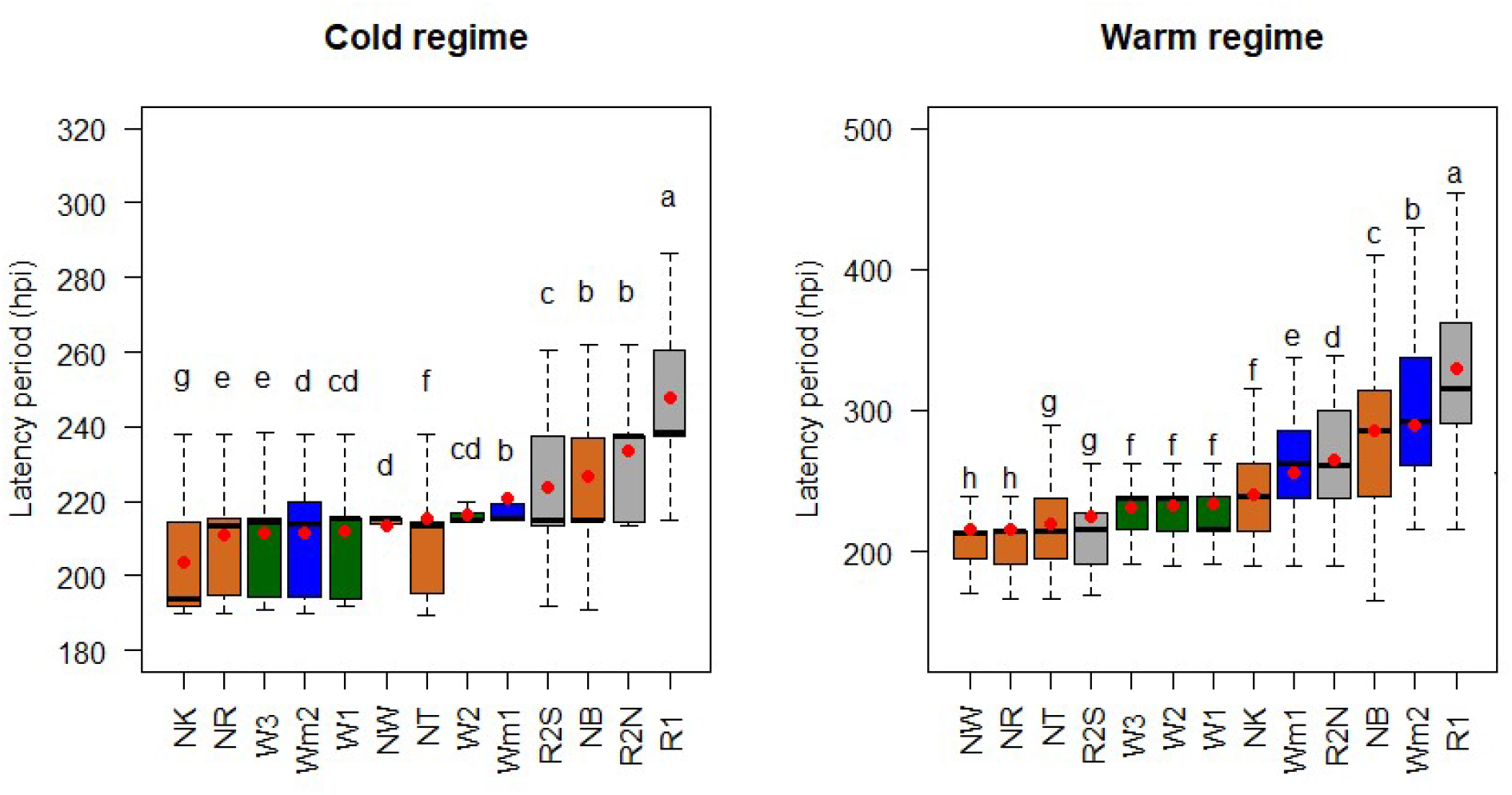
Mean latency period (LP) expressed in hours post-inoculation (hpi) and variability of the LP for the 13 *Puccinia striiformis* f. sp. *tritici* isolates tested under a cold (15°C-10°C) or a warm (25°C-16°C) regime. Letters indicate significant differences (*p*-value <0.05; Kruskal-Wallis test). The red dots indicate the mean values. The races currently most common in Europe are indicated by the color of the boxes: green for ‘Warrior’, blue for ‘Warrior(-)’ and brown for the “new” isolates. The old reference isolates are represented by gray boxes.

**Figure 6.**
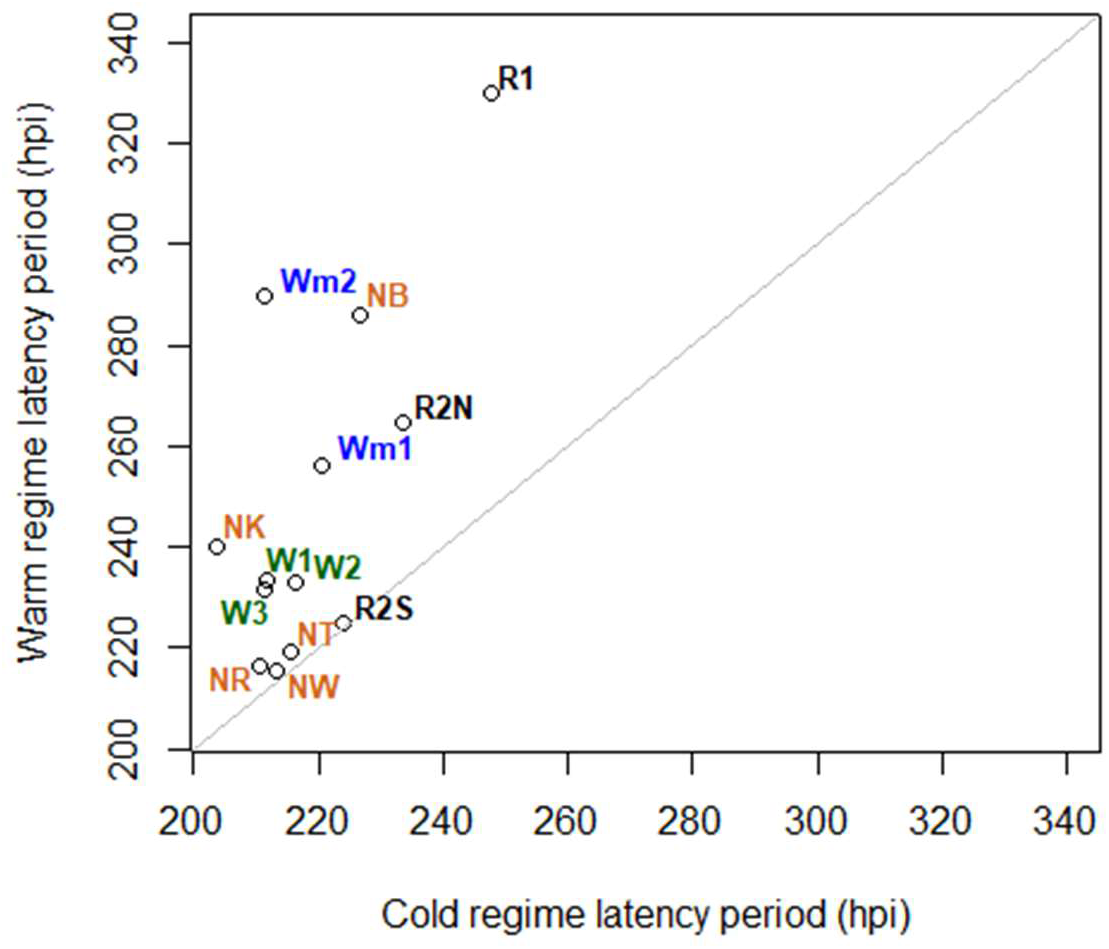
Differences in the latency periods (LPs) of the 13 *Puccinia striiformis* f. sp. *tritici* isolates under the cold (15°C-10°C) and warm (25°C-16°C) regimes. LP is expressed in hours post inoculation (hpi). The isolates above the bisector had a longer LP under the warm regime than under the cold regime. The closer to the bisector the isolate lies, the more similar its LP values under the two thermal regimes. The races currently most common in Europe are indicated by the color code: green for ‘Warrior’, blue for ‘Warrior(-)’ and brown for the “new” isolates. The old reference isolates are represented in black.

NR, a recent isolate from the new British race ‘Red 24/Warrior(-)’, was among the best-performing isolates (short LP) under both temperature regimes. By contrast, NB, from the recent British race ‘Blue 7/Solstice’, had a long LP under both regimes and appeared to be worst performing of the recent isolates. Both isolates from the ‘Warrior(-)’ race (Wm1 and Wm2) had long LPs under the warm regime, whereas all of the isolates from the ‘Warrior’ race (W3, W1 and W2) had a similar, intermediate LP under both regimes. The NT isolate of the ‘Triticale’ race was one of the best-performing isolates under the warm regime, whereas it had an intermediate LP under the cold regime. By contrast, the NK isolate from the ‘Kranich’ race had a short LP under the cold regime and an intermediate LP under the warm regime.

A factor map, projecting the LPs under both the cold and warm regimes (Figure S1), identified three groups, (i) a group of the most aggressive isolates, (ii) a group of less aggressive isolates including PstS0 (R2N and NB) and PstS10 (Wm1 and Wm2), and (iii) the least aggressive isolate, R1 (PstS0).

Isolates with similar, low LPs under the two regimes are highlighted in Figure 6: NR, NW, NT and R2S, i.e. the new British ‘Red 24/Warrior(-)’ race, the ‘Warrior/Kranich’ French race, the ‘Triticale’ race and the southern French reference race, respectively. Isolates from the ‘Warrior’ race (W1, W2 and W3) had an intermediate LP at both regimes that was shorter than that of ‘Warrior(-)’ isolates, and they formed a single group. Interestingly, ‘Warrior(-)’ isolates (Wm1 and Wm2), the ‘Blue 7/Solstice’ isolate (NB) and the northern reference isolates (R1 and R2N) had a longer LP under both regimes, with R1, the oldest and least virulent isolate, having the longest LP. The differential behavior of “new” European *Pst* isolates under the cold and warm regimes is highlighted in Figure 6: all these isolates are positioned above the bisector line, indicating better adaptation to the warm regime. The distance to the bisector line ranged from 20 to 70 hpi, indicating a potential gain of 1 to 3 days under warm conditions relative to “old” isolates.

### Relationship between the aggressiveness components IE and LP

Principal component analysis (PCA) was performed to visualize the behavior of the 13 *Pst* isolates in terms of IE and LP under various thermal regimes (LP under cold (15°C/10°C) and warm (25°C/16°C) regimes, and IE at 5°C, 10°C, 15°C and 20°C). We defined six groups based on coordinates on the first two PCA axes, accounting for 59.9% of the variability (Figure 7). IE at 5°C was the variable least represented in the factorial plane. PC1 and PC2 were positively correlated with LP under both the cold and warm regimes, and PC1 was negatively correlated with IE at 10°C (Table S2). LP under both the cold and warm regimes was inversely correlated with IE at 10°C (Table S2), highlighting the existence of isolates with a high IE at warm temperatures and isolates with a low IE at warm temperatures. Two groups (1 and 6; Figure 7) consisted of single isolates and behaved as outliers: NT, characterized by a high IE at warm temperatures and an overall short LP, and R1, characterized by a low IE at cold temperatures and an overall long LP. Groups 2, 3 and 4 consisted of isolates with a short LP, but NW and NR had a higher IE at 10°C, whereas isolates from groups 3 and 4 had more similar values of IE. Isolates from group 5 (Wm1, R2N and NB) had low IE and long LP values, and were considered to be the least aggressive isolates overall.

**Figure 7.**
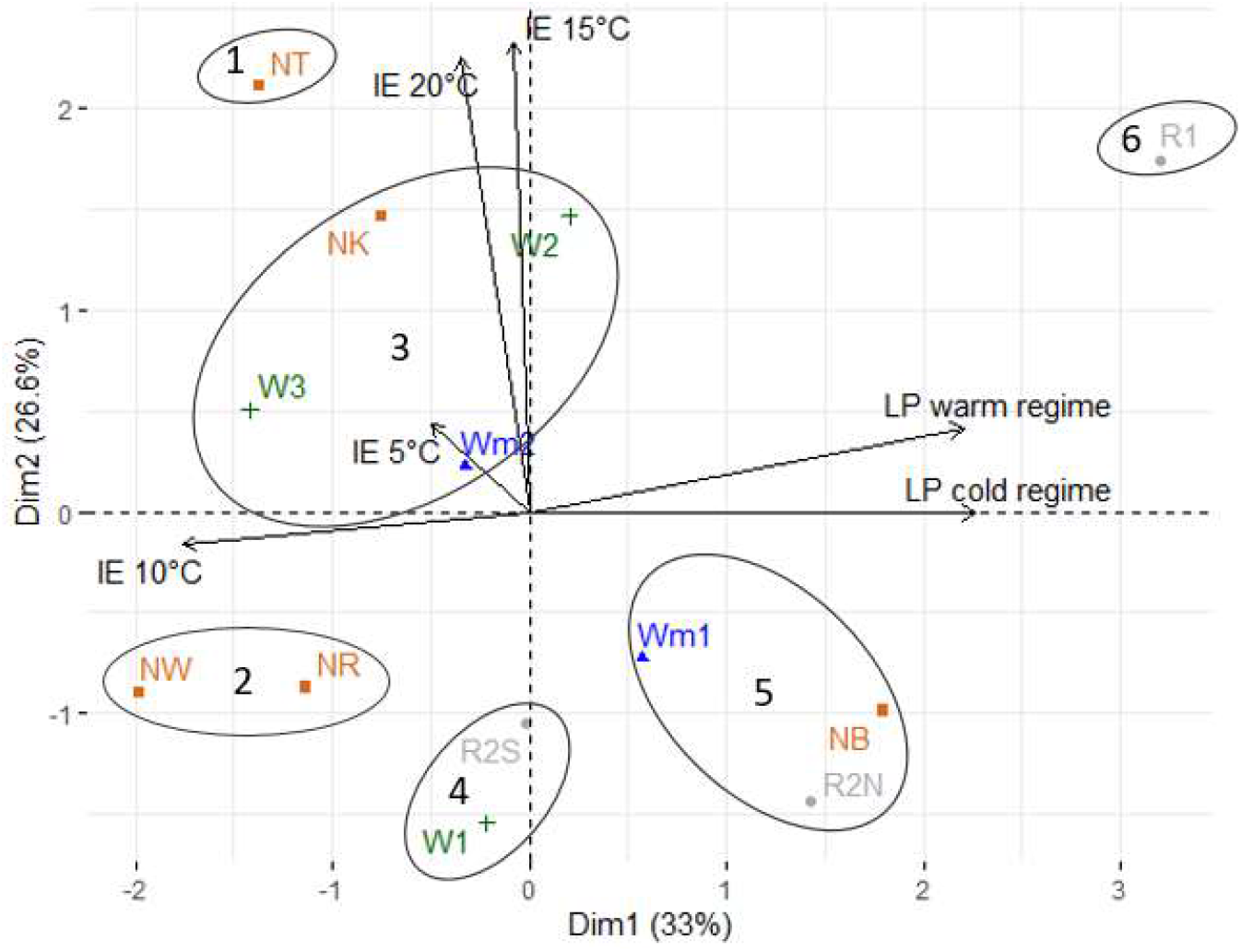
Factor map (PCA) plot of the 13 isolates of *Puccinia striiformis* f. sp. *tritici* projected onto the first two principal components according to estimated infection efficiency (IE) and latent period (LP) under different thermal regimes (LP under the cold (15°C/20°C) and warm (25°C/16°C) regimes, and IE at 5°C, 10°C, 15°C and 20°C).

## Discussion

### Differences in thermal aptitude can help to explain the establishment of new pathogen lineages

Under changing climatic conditions, with an overall warming of the European continent, the emergence of several diseases – sporadic incursions of plant pathogens but also their establishment in large spatial areas – have been reported in recent years. New lineages of pathogens already widespread in Europe, the re-emergence of pathogen species after many years of absence, and the identification of new pathogen species all represent significant threats to wheat production. From 2011 onwards, the exotic *Pst* race Warrior rapidly colonized European wheat areas, replacing older European races (Hovmøller *et al*., 2016). *P. graminis f. sp. tritici,* the causal agent of stem rust, which had been barely present in Europe for decades, made several incursions at the beginning of the 2020s (Patpour *et al*., 2022), while wheat blast, caused by *Pyricularia oryzae*, has caused damage in Asia and Africa over the last few years and may soon arrive at the gates of Europe (Latorre *et al*., 2023). Several previous studies have highlighted the diversity of thermal aptitude and its potential impact on the emergence of new *Pst* races in Europe, including France (Mboup *et al*., 2012; de Vallavieille-Pope *et al*. in 2018; Vidal *et al*. 2022), in Middle Eastern and Mediterranean areas (El Amil *et al*., 2022), and in North America (Milus *et al*., 2009; Lyon & Broders, 2017). In this epidemiological context, our experimental results provide relevant complementary information — with elements of explanation but also signs of complexity — about thermal aptitude, which is thought to be one of the factors underlying the success of new emerging races of *Pst* in Europe. Variations in thermal aptitude are easier to characterize for pathogens that are already established, as sufficiently diverse fungal material must be available for analysis, which may be a challenging requirement for populations dominated by clonal lineages like those of *Pst*. The 2000-2020 period was particularly favourable for such analyses, as several successive replacements of *Pst* races occurred over this period.

For all the isolates tested here, infection efficiency (IE) was highest for incubation at 5°C and the latency period (LP) was shortest under the cold regime (15°C-20°C). This result is consistent with the already established preference of *Pst* for cooler climates. All isolates were highly sensitive to temperature, both during the infection period, including spore germination, and during the latency period (i.e. the days preceding the sporulation), as demonstrated by the large differences in IE (from 0.3 to 26%) and in LP (from 220 to 330 hpi), respectively. The significance of the differences in IE and LP between thermal regimes is important, but difficult to interpret at the level of individual isolates. This finding highlights the ability of European *Pst* populations to adapt to major changes in temperature with an impact on their aggressiveness. The most salient results of this study (Figures 3 and 5) are the differences in thermal behavior (i) between the “old” reference isolates (R2S, R2N and R1) and the two most recent dominant European lineages, ‘Warrior’ and ‘Warrior(-)’, and (ii) between the ‘Warrior’ and ‘Warrior(-)’ lineages.

The reference isolate from southern France was theoretically better adapted to warm conditions, but did not have the necessary virulence genes to develop on current wheat varieties (de Vallavieille-Pope *et al*., 2018). LP analysis also revealed that some other “new” isolates, such as those of the ‘Triticale’ (NT) and ‘Warrior/Kranich’ (NK) races, were particularly aggressive, especially under the warm regime, with no negative impact of changes in temperature. However, such isolates have been and remain relatively uncommon in European wheat-growing areas. One of the most recent isolates, from the new British ‘Red 24/Warrior(-)’ (NR) race, appeared to be particularly aggressive, with a short LP under both the warm and cold regimes, especially relative to the ‘Blue 7/Solstice’ (NB) race, which had a longer LP (Figures 5 and 6). This aggressiveness, together with a complex virulence combination, according to observations made by the NIAB (UK CPVS Annual report, 2017), suggests that the ‘Red 24/Warrior(-)’ (NR) race has a high epidemic potential.

### Differences in thermal aptitude between isolates for a given aggressiveness component – IE or LP – suggest an impact of geographic origin within the new ‘Warrior’ and ‘Warrior(-)’ races

Certain differences in IE highlighted thermal adaptation within the same race, depending on the geographic origin of the isolates. This type of difference first appeared in comparisons of the French reference isolates: the “inversions” of performance rankings between 5°C and 10°C for the southern and northern French reference isolates (R2S, R2N, respectively; Figure 4) are consistent with the evidence of thermal adaptation reported by Mboup *et al*. (2012). This finding provides support for the reliability of our results for the thermal behavior of *Pst* races, indicating that the differences in IE probably reflect real differences in thermal performance between isolates. Within the ‘Warrior’ and ‘Warrior(-)’ races, we highlighted similar differences in IE constituting signs of thermal adaptation, according to the geographic origin of the isolates, with the two French isolates having an advantage at higher temperatures – starting from 10°C for ‘Warrior’ and from 15°C for ‘Warrior(-)’ – over strains from the more northerly areas of the UK and Denmark (Table 2). Given the small number of isolates tested, some caution is required in interpretation, but this conclusion is supported by the consistent results repeatedly obtained in this study.

Significant differences in LP were also found between isolates. Contrary to the findings of other experiments performed with the same method and common isolates (de Vallavieille-Pope *et al*., 2018), LP appeared to be shorter under the cold regime than under the warm regime. This difference may be explained by differences in performance of the technical facilities used in the past, and justifies the use of reference isolates and caution concerning the precision of the results that can be obtained with this type of experiment. However, the robustness of the results obtained here is supported by the longer LP under the cold regime and the shorter LP under the warm regime of the southern French reference isolate (R2S) relative to the other isolates, consistent with the results obtained by Mboup *et al*. (2012). Furthermore, a recent reanalysis of a published dataset (de Vallavieille-Pope *et al*., 2018) by Vidal *et al*. (2022) suggested an optimal temperature of 15°C to 20°C and highlighted the existence of few differences in LP at temperatures between 7°C and 20°C, indicating a more limited variability of thermal aptitude for LP than for IE.

### The variability of the temperature response of aggressiveness components does not explain the replacement of ‘Warrior’ by ‘Warrior(-)’: a result to be interpreted with caution

The isolates from the ‘Warrior’ and ‘Warrior(-)’ races did not appear to be more aggressive than the isolates of the other races tested. Instead, they displayed “generalist” behavior in terms of their thermal requirements, particularly for the LP of ‘Warrior’ isolates. W1, W2 and W3 performed equally well under the various thermal regimes tested. However, beyond this generalist behavior, differences emerged between ‘Warrior’ and ‘Warrior(-)’, depending on the aggressiveness trait considered. ‘Warrior’ appeared to be less well adapted to cold conditions than ‘Warrior(-)’ according to the IE values obtained, suggesting that its thermal aptitude resembles that of the southern French reference isolate (RS). Conversely, ‘Warrior(-)’ appeared to be less well adapted to warm conditions than ‘Warrior’ according to the LP values obtained, suggesting that its thermal aptitude resembles that of the northern French reference isolate (RN). This difference highlights the complexity of the analysis and of drawing conclusions about the epidemiological success of one group of isolates relative to another, as the complementarity between aggressiveness traits must be taken into account. Similar observations have been made for the reference isolates R1 and R2N collected in northern areas, as mentioned above, all part of the PstS0 genetic group prevalent in Europe before the first ‘Warrior’ incursion, and considered to be a group best suited to colder climates (Mboup *et al*., 2012).

Results for large temporal and spatial scales should be interpreted with caution, taking other more powerful adaptive dynamics into account. A fitness cost of thermal aptitude or a more advantageous virulence profile relative to the resistances deployed in the landscape following the acquisition of virulence genes by new lineages probably accounts for the difference in competitive success between the ‘Warrior’ and ‘Warrior(-)’ races. Caution is particularly important when interpreting the results for *Pst*, which has clonal lineages and in which adaptations to different factors result from migration and cannot easily accumulate over time due to the local absence or very low levels of genetic exchange between lineages. However, this lineage homogeneity is tempered by high levels of variability in thermal aptitude between ‘Warrior’ isolates, as shown by Pope et al. (2018). The successful ‘Warrior’ and ‘Warrior(-)’ races seem better adapted to cold than to warm conditions in terms of LP (as shown in Figure 6), contrary to the expected trend towards adaptation and greater tolerance of warm conditions in response to climate change. Ali *et al*. (2017) initially attributed the rapid colonization of European wheat-growing areas by ‘Warrior’ in recent years to a better adaptation of this race to warmer climates, although this race was subsequently replaced by ‘‘Warrior(-)’, especially in France (https://agro.au.dk/forskning/internationale-platforme/wheatrust/yellow-rust-tools-maps-and-charts/races-changes-across-years). The greater success of Warrior than of other exotic races (e.g. the PstS2 group which was widely prevalent in Asia and Africa; Ali *et al*., 2017) in Europe in the recent past can also be explained by the very small proportion of the wheat-growing area (e.g. 15% in France; Vidal *et al*., 2022) displaying a low risk of infection with this *Pst* race before its emergence. The interaction between virulence spectrum and thermal aptitude was investigated in detail by comparing the behavior of ‘Warrior’ (PstS7) and PstS2, which had a very limited impact in France despite being better adapted to warm conditions (de Vallavieille-Pope *et al*., 2018; Vidal *et al*., 2022). The subsequent success of ‘Warrior(-)’, and of new variants, was conferred by a virulence gene enabling these races to attack wheat varieties containing an as yet unidentified resistance gene.

Comparisons of thermal aptitude between the three ‘Warrior’ isolates and the two ‘Warrior (-)’ isolates are relevant, but should be interpreted with caution, given the small sample size. We observed significant differences between isolates, but it is not possible to generalize our data to the particular races or geographic origins of the isolates studied here (France, UK, and Denmark), as the isolates studied are the only representatives of these categories available and may not be truly representative. For the conclusion that a race is more adapted to warm or cold temperatures to be draw, sufficient numbers of isolates from the race concerned must be compared with sufficient isolates from another race. Indeed, it is not rare to observe significant differences between isolates of the same race, as demonstrated by Vallavieille-Pope *et al*. (2018) for ‘Warrior’ and by Fontyn *et al*. (2022) for *P. triticina*. Populations adapt continually to different factors, including climatic conditions, leading to an increase in pathogen fitness and aggressiveness not necessarily associated with the presence or absence of virulence genes. Furthermore, experimental results obtained with seedlings can provide general clues concerning adaptation, but may not entirely explain the population dynamics of field-grown plants, particularly at the adult stage.

It is essential to monitor the thermal aptitude of new *Pst* lineages, in addition to their virulence spectrum. If we are to improve our understanding of the influence of the thermal aptitude of these lineages on their success in Europe, in prospective or retrospective studies, we will need to improve the protocols for estimating aggressiveness traits. For instance, new protocols have been proposed for the measurement of IE in *P. triticina* based on single-spore isolation (Fontyn *et al*., 2022), but the numbers of isolates and plants studied here were too great for this protocol to be used. Modification of the range of temperatures used for testing may also be required. Previous studies have suggested an optimum temperature of about 10°C for plant infection (Mboup *et al*., 2012), subsequently refined to about 8°C for the PstS0, PstS2 and PstS7 genetic groups on the susceptible cv Victo (de Vallavieille-Pope *et al*., 2018). The determination of a more precise optimum temperature would require experimentation at lower temperatures but this would be difficult given the thermal requirements for wheat growth and technical limitations. It would also be useful to characterize other aggressiveness components in *Pst*, such as sporulation capacity (the number of spores produced per lesion), which might counterbalance lower competitiveness for another trait.

### The independence of thermal aptitude for IE and LP suggests that temperature may have different consequences at different epidemic stages, early or late in the season

The competitive advantage of a high IE under cold conditions was observed at 5°C in ‘Warrior’ isolates (Figure 3), and at 10°C across all profiles, with an overall inverse correlation with LP in both warm and cold conditions (Figure 7). A well-defined ‘Warrior’ group emerged (Figure 6), with an LP shorter than that for ‘Warrior(-)’ under the warm regime, revealing “generalist” behavior. The advantage conferred by this aggressiveness component must be considered in the context of epidemics, and the ways in which each trait influence the dynamics of the epidemic. The relative influence of IE and LP may vary between epidemic stages and be affected by other variables, including climatic factors. For instance, a high level of ability to infect wheat tissues and to persist in these tissues (through a high IE) in cold conditions would make it possible for sporulation to occur at the start of spring. This might be a greater advantage than having a shortener cycle (short LP) at the end of the epidemic season, when conditions are warmer. An isolate could therefore be considered to be favored if it displays a slight advantage for a trait that is limiting under current thermal conditions (e.g. infecting and surviving in cold weather). A competitive disadvantage for a trait that is not limiting under current thermal conditions may be less detrimental (e.g. multiplying faster when it is warmer). This view is consistent with the observed advantage of the short LP in cold conditions of the ‘Warrior’ and ‘Warrior(-)’ races relative to the “old” French reference races (R2S and R2N). The hypothesis that a high IE in cold conditions is of greater advantage than a short LP in warm conditions is highly debatable, but consistent with the results of modelling experiments suggesting that ‘Warrior’ is more competitive against other European races than PstS2 (Vidal *et al*., 2022).

In conclusion, this study provides insight into the potential effects of temperature on the behavior and adaptability of *Pst*, highlighting the importance of considering both geographic origin and the epidemic stage at which certain components of aggressiveness may be more important than others in studies of plant-pathogen interactions in the context of global warming. It shows that, while thermal aptitude is important, it was not the major driver of the success of ‘Warrior(-)’ and the races that succeeded it after 2015, such as ‘Triticale’ and ‘Kranich’.

## Acknowledgments

We thank Laurent Gérard and Nathalie Retout (INRAE BIOGER) for their technical assistance in preparing the experiments. We also thank Chris K Sørensen (Aarhus University, Denmark) and Amelia Hubbard (NIAB, United-Kingdom) for providing study isolates. We are grateful to Claude de Vallavieille-Pope, who retired just before the set-up of the RUSTWATCH project, for having spearheaded the theoretical concepts and experimental approaches used in this study. We thank Julie Sappa for her editorial advice on English usage.

## Author Contributions

ML conceived the original idea and designed the study, with input from FS and HG. HG and FS supervised the funding and administration of the research program with input from ML. ML and KM planned and performed the experiments with input from TV. KM and TV performed the data analysis. All the authors contributed to the interpretation of the results. KM, FS and HG wrote the manuscript, with input from TV. All authors provided critical feedback and approved the final version of the manuscript.

## Supplementary material

**Table S1.**
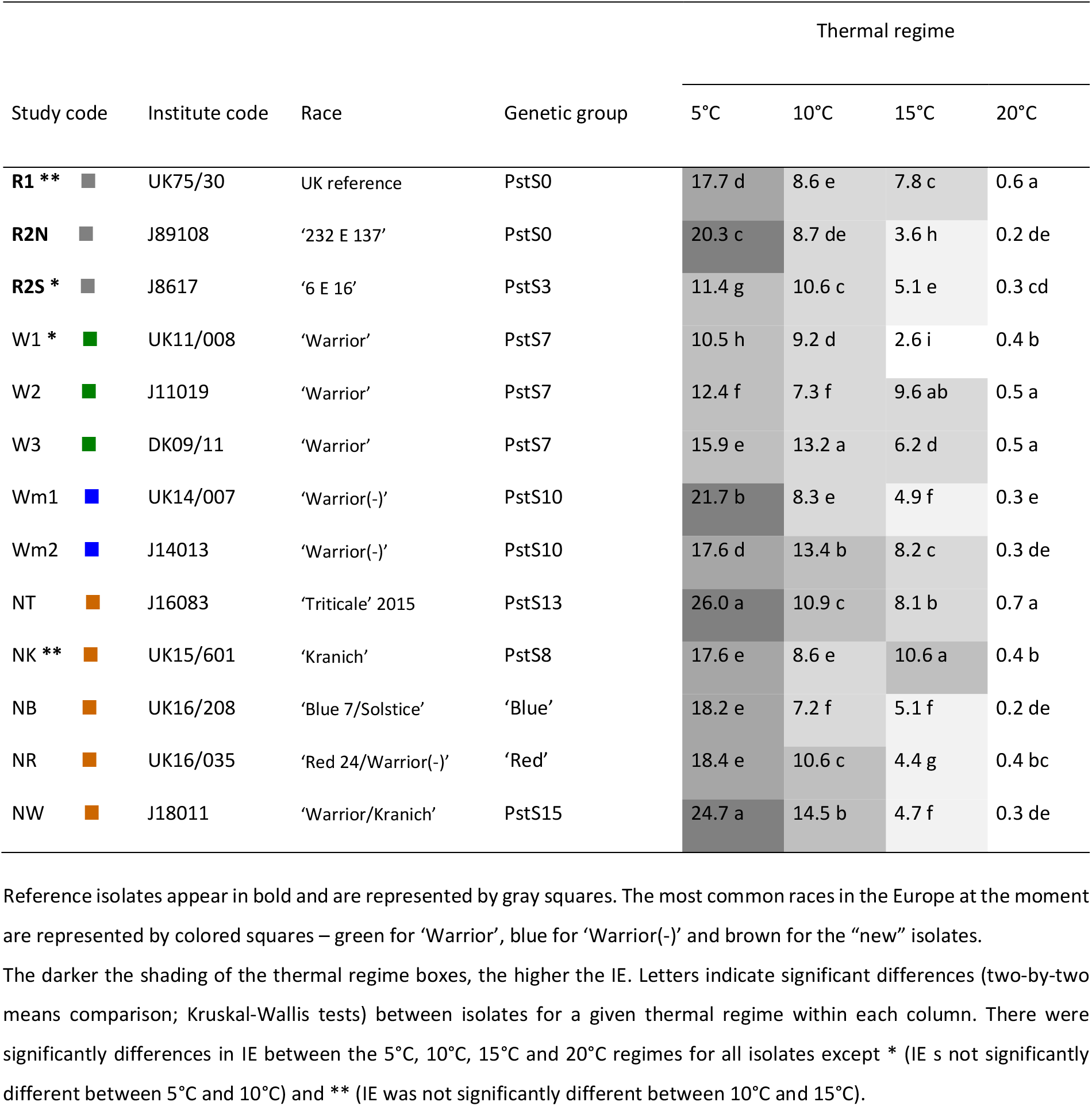
Mean infection efficiency (IE) for the 13 *Puccinia striiformis* f. sp. *tritici* isolates tested under four thermal regimes (5°C, 10°C, 15°C, and 20°C) during the first 24 hours after inoculation

| Study code | Institute code | Race | Genetic group | Thermal regime |  |  |  |
| --- | --- | --- | --- | --- | --- | --- | --- |
|  |  |  |  | 5°C | 10°C | 15°C | 20°C |
| **R1 \*\*** | UK75/30 | UK reference | PstS0 | 17.7 d | 8.6 e | 7.8 c | 0.6 a |
| **R2N** | J89108 | '232 E 137' | PstS0 | 20.3 c | 8.7 de | 3.6 h | 0.2 de |
| **R2S \*** | J8617 | '6 E 16' | PstS3 | 11.4 g | 10.6 c | 5.1 e | 0.3 cd |
| W1 * | UK11/008 | 'Warrior' | PstS7 | 10.5 h | 9.2 d | 2.6 i | 0.4 b |
| W2 | J11019 | 'Warrior' | PstS7 | 12.4 f | 7.3 f | 9.6 ab | 0.5 a |
| W3 | DK09/11 | 'Warrior' | PstS7 | 15.9 e | 13.2 a | 6.2 d | 0.5 a |
| Wm1 | UK14/007 | 'Warrior(-)' | PstS10 | 21.7 b | 8.3 e | 4.9 f | 0.3 e |
| Wm2 | J14013 | 'Warrior(-)' | PstS10 | 17.6 d | 13.4 b | 8.2 c | 0.3 de |
| NT | J16083 | 'Triticale' 2015 | PstS13 | 26.0 a | 10.9 c | 8.1 b | 0.7 a |
| NK ** | UK15/601 | 'Kranich' | PstS8 | 17.6 e | 8.6 e | 10.6 a | 0.4 b |
| NB | UK16/208 | 'Blue 7/Solstice' | 'Blue' | 18.2 e | 7.2 f | 5.1 f | 0.2 de |
| NR | UK16/035 | 'Red 24/Warrior(-)' | 'Red' | 18.4 e | 10.6 c | 4.4 g | 0.4 bc |
| NW | J18011 | 'Warrior/Kranich' | PstS15 | 24.7 a | 14.5 b | 4.7 f | 0.3 de |

Reference isolates appear in bold and are represented by gray squares. The most common races in the Europe at the moment are represented by colored squares – green for 'Warrior', blue for 'Warrior(-)' and brown for the "new" isolates.
The darker the shading of the thermal regime boxes, the higher the IE. Letters indicate significant differences (two-by-two means comparison; Kruskal-Wallis tests) between isolates for a given thermal regime within each column. There were significantly differences in IE between the 5°C, 10°C, 15°C and 20°C regimes for all isolates except \* (IE is not significantly different between 5°C and 10°C) and \*\* (IE was not significantly different between 10°C and 15°C).

**Table S2.** Coefficient of the correlation between the first two principal components (PC1, PC2) according to infection efficiency (IE) and latent period (LP), estimated for the 13 *Puccinia striiformis* f. sp. *tritici* isolates under the various thermal regimes (LP under cold (15°C/20°C) and warm (25°C/16°C) regimes, and IE at 5°C, 10°C, 15°C and 20°C).

|  | PC1 | PC2 |
| --- | --- | --- |
| IE 5°C | -0.1924 | 0.1682 |
| IE 10°C | -0.6755 | -0.0621 |
| IE 15°C | -0.0305 | 0.8930 |
| IE 20°C | -0.1343 | 0.8619 |
| LP cold regime | 0.8673 | -0.0013 |
| LP warm regime | 0.8457 | 0.1591 |
| Eigenvalue | 1.98 | 1.60 |
| Percentage of variance | 33.00 | 26.63 |

**Figure S1.**
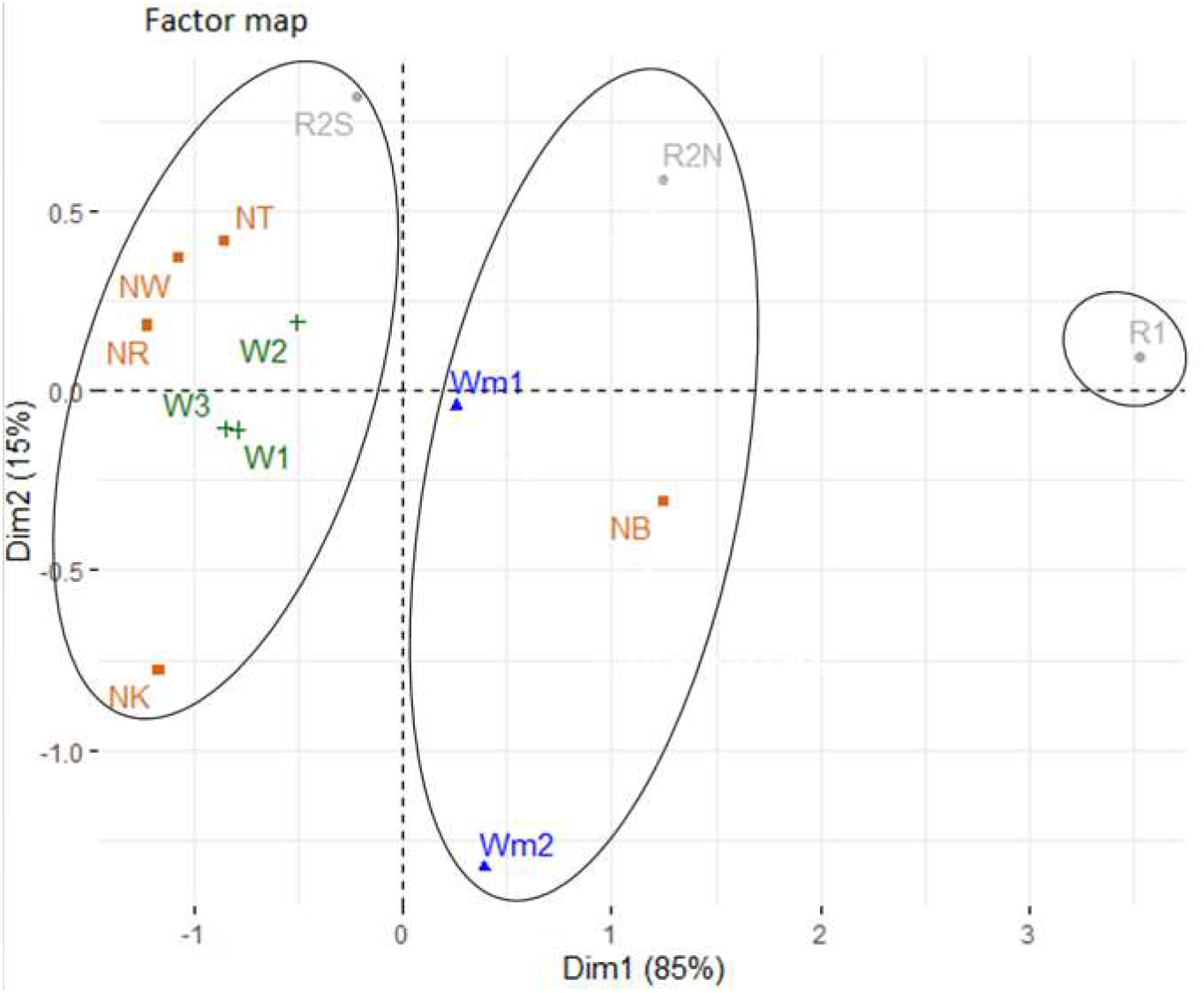
Factor map (PCA) plot for the 13 isolates of *Puccinia striiformis* f. sp. *tritici* projected onto the first two principal components according to LP under the cold regime and LP under the warm regime. The races currently most common in Europe are shown in color – green for ‘Warrior’, blue for ‘Warrior(-)’ and brown for the “new” isolates. The old reference isolates are represented in gray.

## References

Ali S, Rodriguez-Algaba J, Thach T, Sørensen CK, Hansen JG, Lassen P, Nazari K, Hodson DP, Justesen AF, Hovmøller MS (2017) Yellow rust epidemics worldwide were caused by pathogen races from divergent genetic lineages. Frontiers in Plant Science 8: 1057. https://doi.org/10.3389/fpls.2017.01057

Andrade C, Leite SM, Santos JA (2012) Temperature extremes in Europe: overview of their driving atmospheric patterns. Natural Hazards and Earth System Sciences 12: 1671–1691. https://doi.org/10.5194/nhess-12-1671-2012

Andrivon D, Pilet F, Montarry J, Hafidi M, Corbière R, Achbani el H, Pellé R, Ellissèche D (2007) Adaptation of *Phytophthora infestans* to partial resistance in potato: evidence from French and Moroccan populations. Phytopathology, 97: 338–343. https://doi.org/10.1094/PHYTO-97-3-0338

Boshoff WHP, Pretorius ZA, van Niekerk BD (2002) Establishment, distribution, and pathogenicity of *Puccinia striiformis* f. sp. *tritici* in South Africa. Plant Disease, 86: 485–492. https://doi.org/10.1094/PDIS.2002.86.5.485

Boixel AL, Chelle M, Suffert F (2022) Patterns of thermal adaptation in a globally distributed plant pathogen: local diversity and plasticity reveal two-tier dynamics. Ecology and Evolution, 12: e8515. https://doi.org/10.1002/ece3.8515

Chen F, Duan G-H, Li D-L, Zhan J (2017) Host resistance and temperature-dependent evolution of aggressiveness in the plant pathogen *Zymoseptoria tritici*. Frontiers in Microbiology, 8: 1217. https://doi.org/10.3389/fmicb.2017.01217

Chen W, Zhang Z, Chen XM, Meng Y, Huang L, Kang Z, Zhao J (2021) Field production, germinability, and survival of *Puccinia striiformis* f. sp. *tritici* teliospores in China. Plant Disease, 105: 2122–2128. https://doi.org/10.1094/PDIS-09-20-2018-RE

De Mendiburu F, Yaseen M (2020). agricolae: Statistical Procedures for Agricultural Research. R package version 1.4.0. https://myaseen208.com/agricolae/

Dennis, JI (1987) Temperature and wet-period conditions for infection by *Puccinia striiformis* f. sp*. tritici* race 104e137a+. Transactions of the British Mycological Society, 88: 119–121. https://doi.org/10.1007/BF02879166

Douville H, Raghavan K, Renwick J, Allan RP, Arias PA, Barlow M, Cerezo-Mota R, Cherchi A, Gan TY, Gergis J, Jiang D, Khan A, Pokam Mba W, Rosenfeld D, Tierney J, Zolina O (2021) Water cycle changes. Contribution of Working Group I to the Sixth Assessment Report of the Intergovernmental Panel on Climate Change, Chapter 8. https://doi.org/10.1017/9781009157896.010

El Amil R, Shykoff JA, Vidal T, Boixel AL, Leconte M, Hovmøller MS, Nazari K, de Vallavieille-Pope C (2022) Diversity of thermal aptitude of Middle Eastern and Mediterranean *Puccinia striiformis* f. sp. *tritici* isolates from different altitude zones. Plant Pathology, 71: 1674–1687. https://doi.org/10.1111/ppa.13613

Figueroa M, Dodds PN, Henningsen EC (2020) Evolution of virulence in rust fungi - Multiple solutions to one problem. *Current Opinion in Plant Biology*, Biotic interactions, 56: 20–27. https://doi.org/10.1016/j.pbi.2020.02.007

Fontyn C, Meyer KJG, Boixel A-L, Delestre G, Piaget E, Picard C, Suffert F, Marcel TC, Goyeau H (2023) Evolution within a given virulence phenotype (pathotype) is driven by changes in aggressiveness: a case study of French wheat leaf rust populations. Peer Community Journal, e39. https://doi.org/10.24072/pcjournal.264

Fontyn C, Zippert A-C, Delestre G, Marcel TC, Suffert F, Goyeau H (2022). Is virulence phenotype evolution driven exclusively by *Lr* gene deployment in French *Puccinia triticina* populations? Plant Pathology, 71: 1511–1524. https://doi.org/10.1111/ppa.13599

Garrett KA, Dendy SP, Frank EE, Rouse MN, Travers SE (2006) Climate change effects on plant disease: genomes to ecosystems. Annual Review of Phytopathology 44, 1: 489–509. https://doi.org/10.1146/annurev.phyto.44.070505.143420

Gulev SK, Thorne PW, Ahn J, Dentener FJ, Domingues CM, Gerland S, Gong D, Kaufman DS, Nnamchi HC, Quaas J, Rivera JA, Sathyendranath S, Smith SL, Trewin B, von Schuckmann K, Vose RS (2021) Changing state of the climate system. Contribution of Working Group I to the Sixth Assessment Report of the Intergovernmental Panel on Climate Change, Chapter 2. https://doi.org/10.1017/9781009157896.004

Helfer S (2014) Rust fungi and global change. New Phytologist, 201: 770–80. https://doi.org/10.1111/nph.12570

Hovmøller MS, Walter S, Bayles RA, Hubbard A, Flath K, Sommerfeldt N, Leconte M, Czembor P, Rodriguez-Algaba J, Thach T, Hansen JG, Lassen P, Justesen AF, Ali S, de Vallavieille-Pope C (2016) Replacement of the European wheat yellow rust population by new races from the centre of diversity in the near-Himalayan region. Plant Pathology, 65: 402–411. https://doi.org/10.1111/ppa.12433

Hubbard A, Lewis CM, Yoshida K, Ramirez-Gonzalez RH, de Vallavieille-Pope C, Thomas J, Kamoun S, Bayles R, Uauy C, Saunders DGO. (2015) Field pathogenomics reveals the emergence of a diverse wheat yellow rust population. Genome Biology, 16: 23. https://doi.org/10.1186/s13059-015-0590-8

Hubbard A, Wilderspin S, Holdgate S (2017) United Kingdom Cereal Pathogen Virulence Survey 2017 Annual Report. NIAB, Huntingdon Road, Cambridge, CB3 0LE

Juroszek P, Racca P, Link S, Farhumand J, Kleinhenz B (2020) Overview on the review articles published during the past 30 years relating to the potential climate change effects on plant pathogens and crop disease risks. Plant Pathology, 69: 179–93. https://doi.org/10.1111/ppa.13119

Lannou C (2012) Variation and selection of quantitative traits in plant pathogens. Annual Review of Phytopathology, 50: 319–338 https://doi.org/10.1146/annurev-phyto-081211-173031

Latorre SM, Were VM, Foster AJ, Langner T, Malmgren A, Harant A, Asuke S, Reyes-Avila S, Gupta DR, Jensen C, Ma W, Uddin Mahmud N, Shåbab Mehebub M, Mulenga RM, Muzahid ANM, Paul SK, Fajle Rabby SM, Rahat AAM, Ryder L, Shrestha R-K, Sichilima S, Soanes DM, Singh PK, Bentley AR, Saunders DGO, Tosa Y, Croll D, Lamour KH, Islam T, Tembo B, Talbot NJ, Burbano HA, Kamoun S (2023) Genomic surveillance uncovers a pandemic clonal lineage of the wheat blast fungus. Public Library of Science Biology, 21: e3002052. https://doi.org/10.1371/journal.pbio.3002052

Lê S, Josse J, Husson F (2008) FactoMineR: A package for multivariate analysis. Journal of Statistical Software, 25: 1–18. https://www.jstatsoft.org/article/view/v025i01

Lee JY, Marotzke J, Bala G, Cao L, Corti S, Dunne JP, Engelbrecht F, Fischer E, Fyfe JC, Jones C, Maycock A, Mutemi J, Ndiaye O, Panickal S, Zhou T (2021) Future global climate: scenario based projections and near-term information. Contribution of Working Group I to the Sixth Assessment Report of the Intergovernmental Panel on Climate Change, Chapter 4. https://doi.org/https://doi.org/10.1017/9781009157896.006

Lyon B, Broders K (2017) Impact of climate change and race evolution on the epidemiology and ecology of stripe rust in central and eastern USA and Canada. Canadian Journal of Plant Pathology, 39: 385–392. https://doi.org/10.1080/07060661.2017.1368713

Ma L, Qiao J, Kong X, Zou Y, Xu X, Chen XM, Hu X(2015) Effect of low temperature and wheat winter-hardiness on survival of *Puccinia striiformis* f. sp. *tritici* under controlled conditions. Public Library of Science One, 10: e0130691. https://doi.org/10.1371/journal.pone.0130691

Mboup M, Bahri B, Leconte M, de Vallavieille-Pope C, Kaltz O, Enjalbert J (2012) Genetic structure and local adaptation of European wheat yellow rust populations: the role of temperature-specific adaptation. Evolutionary Applications, 5: 341–52. https://doi.org/10.1111/j.1752-4571.2011.00228.x

McDonald BA, Linde C (2002) Pathogen population genetics, evolutionary potential, and durable resistance. Annual Review of Phytopathology, 40: 349–79. https://doi.org/10.1146/annurev.phyto.40.120501.101443

Milus EA, Kristensen K, Hovmøller MS (2009) Evidence for increased aggressiveness in a recent widespread strain of *Puccinia striiformis* f. sp. *tritici* causing yellow rust of wheat. Phytopathology, 99: 89–94. https://doi.org/10.1094/PHYTO-99-1-0089

Milus EA, Seyran E, McNew R (2006) Aggressiveness of *Puccinia striiformis* f. sp. *tritici* isolates in the South-Central United States. Plant Disease, 90: 847–852. https://doi.org/10.1094/PD-90-0847

Novotná M, Hloucalová P, Skládanka J, Pokorný R (2017) Effect of weather on the occurrence of *Puccinia graminis* subsp. *graminicola* and *Puccinia coronata* f. sp. *lolii* at *Lolium perenne* L. and *Deschampsia caespitosa* (L.). Acta Universitatis Agriculturae et Silviculturae Mendelianae Brunensis, 65: 125–134. https://doi.org/10.11118/actaun201765010125

Pariaud B, Ravigné V, Halkett F, Goyeau H, Carlier J, Lannou C (2009) Aggressiveness and its role in the adaptation of plant pathogens. Plant Pathology, 58: 409–424. https://doi.org/10.1111/j.1365-3059.2009.02039.x

Patpour M, Hovmøller MS, Rodriguez-Algaba J, Randazzo B, Villegas D, Shamanin VP, Berlin A, Flath K, Czembor P, Hanzalova A, Sliková S, Skolotneva ES, Jin Y, Szabo L, Meyer KJG, Valade R, Thach T, Hansen JG, Justesen AF (2022) Wheat stem rust back in Europe: diversity, prevalence and impact on host resistance. Frontiers in Plant Science, 13: 882440. https://doi.org/10.3389/fpls.2022.882440

Prank M, Kenaley SC, Bergstrom GC, Acevedo M, Mahowald NM (2019) Climate change impacts the spread potential of wheat stem rust, a significant crop disease. Environmental Research Letters, 14: 124053. https://doi.org/10.1088/1748-9326/ab57de

R Core Team (2022) R: A language and environment for statistical computing. R Foundation for Statistical Computing, Vienna, Austria. https://www.R-project.org/

Rodriguez-Algaba J, Sørensen CK, Labouriau R, Justesen AF, Hovmøller MS (2019) Susceptibility of winter wheat and triticale to yellow rust influenced by complex interactions between vernalisation, temperature, plant growth stage and pathogen race. Agronomy, 10: 13. https://doi.org/10.3390/agronomy10010013

Sørensen CK, Justesen AF, Hovmøller MS (2013) Spontaneous loss of Yr2 avirulence in two lineages of *Puccinia striiformis* did not affect pathogen fitness. Plant Pathology, 62: 19–27. https://doi.org/10.1111/ppa.12147

Suffert F, Ravigné V, Sache I (2015) Seasonal changes drive short-term selection for fitness traits in the wheat pathogen *Zymoseptoria tritici*. Applied and Environmental Microbiology, 81: 6367– 6379 https://doi.org/10.1128/AEM.00529-15

Thrall PH, Burdon JJ (2003) Evolution of virulence in a plant host-pathogen metapopulation. Science, 299: 1735–1737. https://doi.org/10.1126/science.1080070

Vallavieille-Pope C, Ali S, Leconte M, Enjalbert J, Delos M, Rouzet J (2012) Virulence dynamics and regional structuring of *Puccinia striiformis* f. sp. *tritici* in France between 1984 and 2009. Plant Disease, 96: 131–140. https://doi.org/10.1094/PDIS-02-11-0078

Vallavieille-Pope C, Bahri B, Leconte M, Zurfluh O, Belaid Y, Maghrebi E, Huard F, Huber L, Launay M, Bancal MO (2018) Thermal generalist behavior of invasive *Puccinia striiformis* f. sp. *tritici* strains under current and future climate conditions. Plant Pathology, 67: 1307–1320. https://doi.org/10.1111/ppa.12840

Vidal T, Boixel A-L, Maghrebi E, Perronne R, Cheyron P, Enjalbert J, Leconte M, Vallavieille-Pope C (2022) Success and failure of invasive races of plant pathogens: the case of *Puccinia striiformis* f. sp. *tritici* in France. Plant Pathology, 71: 1525–1536. https://doi.org/10.1111/ppa.13581

